# Antimony 3: Extending human-readable model definitions for SBML Level 3 Core and Packages

**DOI:** 10.64898/2026.04.07.717118

**Authors:** Adel Heydarabadipour, Lucian Smith, Joseph L. Hellerstein, Herbert M. Sauro

## Abstract

Antimony is a human-readable language for defining and sharing models developed by the systems biology community. It enables scientists to describe biochemical networks with a simple syntax, while supporting seamless conversion to and from the Systems Biology Markup Language (SBML) community standard. Since Antimony’s original release, both SBML and modeling practices have evolved significantly, creating a need to update Antimony to maintain its standards compliance and practical relevance. In this paper, we introduce Antimony 3, a comprehensive update that formalizes its cumulative improvements and extends its support for SBML Level 3 Core and Flux Balance Constraints (FBC), Distributions, Layout, and Render packages. Antimony 3 enables model specifications that combine kinetic reactions with flux balance analysis, represent uncertainty using probability distributions, add biological context through annotations, and define publication-ready visualizations, all within a unified plain-text format. Antimony 3 is delivered as a lightweight C/C++ library with a stable C API. It is available through official bindings for Python, Julia, and JavaScript/WebAssembly, as well as a cross-platform desktop GUI, which enables straightforward use across scripting environments, desktop applications, and browser-based tools. Antimony 3 is released as open-source software under the BSD 3-Clause License and is available at https://github.com/sys-bio/antimony.

**Author Summary:** Biological models are typically stored in standardized formats that ensure compatibility across different software tools, but these formats rely on verbose, machine-readable syntax that is difficult for humans to write or inspect directly. Antimony addresses this challenge by providing an intuitive, text-based language for defining biological models that can be automatically converted to and from the Systems Biology Markup Language (SBML). Since Antimony’s original release in 2009, the SBML standard and common modeling workflows have expanded significantly. We developed Antimony 3 to support these advances, enabling researchers to write a single human-readable text file that defines reaction networks, constraint-based objectives, uncertainty in parameters and initial conditions, semantic annotations linking to biological databases, and model diagrams. Antimony 3 is provided as open-source software with broad support across computational environments, making it accessible to researchers in a wide range of workflows.

## Introduction

In systems biology, a biological computational model can be described as a set of species and reaction rules that transform species concentrations over time [1]. With the growth in popularity of such models, community standards such as SBML [2] and CellML [3] emerged to enable the exchange of models between different software tools. Over the last two decades, this standardization has led to a rich ecosystem of standard-compatible model simulators, editors, and repositories. The real impact of these community standard, however, has been the ability to preserve and make reproducible thousands of models published in the last 30 to 40 years of research.

Although community standards have largely resolved the model exchange issue, their machine-readable syntax, typically in XML serializations, remains verbose and makes their manual inspection and modification laborious and error-prone. A further drawback is that language models often struggle with the verbose syntax of XML, which imposes limitations on relying on them for biological modeling. To restore human readability without compromising standards compliance, complementary modeling modalities have emerged. Graphical tools such as CellDesigner [4], PathwayDesigner [5], and CopasiUI [6] provide interactive environments that allow modelers to build and edit models visually, and export standards-compliant files for downstream analysis. Despite these advantages, graphical tools often struggle with complex models, as designing interfaces that remain intuitive and performant at scale is inherently challenging. Programmatic libraries, such as libSBML [7] and libCellML [3], offer a different modality: they expose low-level APIs to allow modelers to script every element of a standards-compliant file. This flexibility, however, comes at a cost: efficient use of these libraries requires strong programming skills and the patience to navigate extensive documentation, each posing significant obstacles for domain scientists who may lack the time or coding background to master these interfaces. Last, we note that language models prefer text-based representations rather than programmatic model descriptions.

To reduce the programming barrier, domain-specific languages (DSLs) for modeling, such as PySCeS [8], yaml2sbml [9], and MobsPy [10], have emerged to abstract boilerplate library calls and allow modelers to focus on biological logic rather than low-level implementation details. While these DSLs offer concise, expressive notation for constructing and editing models, they are usually tied to a particular host language or runtime. Antimony breaks that dependency by providing a plain-text, human-readable syntax that round-trips models to and from SBML. In practice, an Antimony model is simply a text file that can be edited with any text editor, thereby providing a streamlined, standards-compliant means of specifying modeling intent.

Version 1 of Antimony [11] introduced a human-readable syntax that allows modelers to write reactions with minimal punctuation (for example, `A -> B; k*A`). This eliminates the need for explicit type declarations and keeps model scripts concise and easy to debug. To enable large models to be assembled from smaller, reusable components, the syntax supports hierarchical modularity, so any model can be instantiated as a submodule with its own namespace and then linked to other modules. Declaring events, explicit rate rules, and assignment rules are among the other widely used modeling features supported in Antimony.

Since Antimony was released in 2009, both the SBML specification and routine modeling workflows have evolved well beyond what the original Antimony syntax could support. SBML has matured to Level 3 Core [12] and is now widely extended via its packages. These advances offer capabilities that could neither be expressed nor preserved by the first Antimony release.

To keep up with evolving SBML capabilities, Antimony has been incrementally upgraded through successive releases. In this paper, we formalize these cumulative improvements as Antimony 3 and detail a set of major enhancements: (i) full support for SBML Level 3 Core alongside the Flux Balance Constraints (FBC) [13], Distributions [14], Layout [15], and Render [16] packages, enabling modelers to use a single script that combines kinetic reactions, FBA objectives, uncertainty distributions, and graphical representations of the model; (ii) an extended syntax that enables inline annotations, on-the-fly rate queries, probabilistic initialization, and algebraic invariants, thereby eliminating the need for auxiliary scripts; and (iii) availability as a lightweight C/C++ core library with official bindings for Python, Julia, and JavaScript/WASM, plus a Qt GUI, enabling seamless model editing in notebooks, desktop applications, and web apps, as well as straightforward integration into larger workflows. A number of tools support the Antimony syntax, including tellurium [17], MakeSBML [18], Iridium [19], and WebIridium [20].

### Design and Implementation

#### Syntax

The Antimony language was originally inspired by Jarnac’s compact notation [21] and was extended to support modular model construction. Its main design philosophy is to keep the language as human-readable and intuitive as possible by avoiding extraneous punctuation, making explicit declarations optional, and using appropriate defaults. These characteristics are advantageous for LLMs as well. A reaction is defined by listing the reactant species, an arrow, and the product species, followed by an optional rate law after a semicolon. As an example, a simple first-order conversion of `S1` to `S2` with rate constant `k1` is written as `S1 -> S2; k1*S1`. This single line implicitly declares `S1` and `S2` as species (they appear as reactant and product of the reaction) and `k1` a parameter since it is referenced in the rate law. Other model elements follow the same concise pattern. A discrete event, for instance, is defined by its trigger condition and the assignment to perform, as in `E0: at time > 3: k4 = 10`, where event `E0` is triggered when simulation time exceeds `3` (in the model’s time units) and sets parameter `k4` to `10`. Species values are assumed to be concentrations, unless explicitly defined to be amounts with a `substanceOnly` directive.

In Antimony 3 (briefly described in [22]), we expanded the grammar to cover more of SBML Level 3 Core together with the Flux Balance Constraints (FBC), Distributions, Layout, and Render packages. To preserve backward compatibility, we layered the new rules on top of the existing grammar. For example, when introducing new features, we retained the no-declaration principle: a single line such as `A1: 0 = S1 + S2` implicitly defines `A1` as an algebraic rule, while `10 < J2 < 50`simultaneously declares `J2` as a parameter and encodes an SBML constraint. Similarly, to avoid extraneous punctuation we extended the dot notation, which was originally used for submodule access, so that `x.mean = 10` assigns a distribution mean to `x` parameter, and `x.position = {50, 40}`sets the layout coordinates of the graphical object representing `x` entity.

#### Architecture

Antimony’s architecture follows a layered design consisting of language bindings, a C API, and a core engine, implemented in C++, that enables lossless, bidirectional translation between the Antimony syntax and SBML format. User artifacts, whether Antimony text or SBML files, are passed through language-specific interfaces, which route all operations through a C API into the core engine. Within the core, models are translated between Antimony and SBML representations. Figure 1 presents a schematic overview of the translation pipeline, and its components are discussed in further detail below.

**Fig 1.**
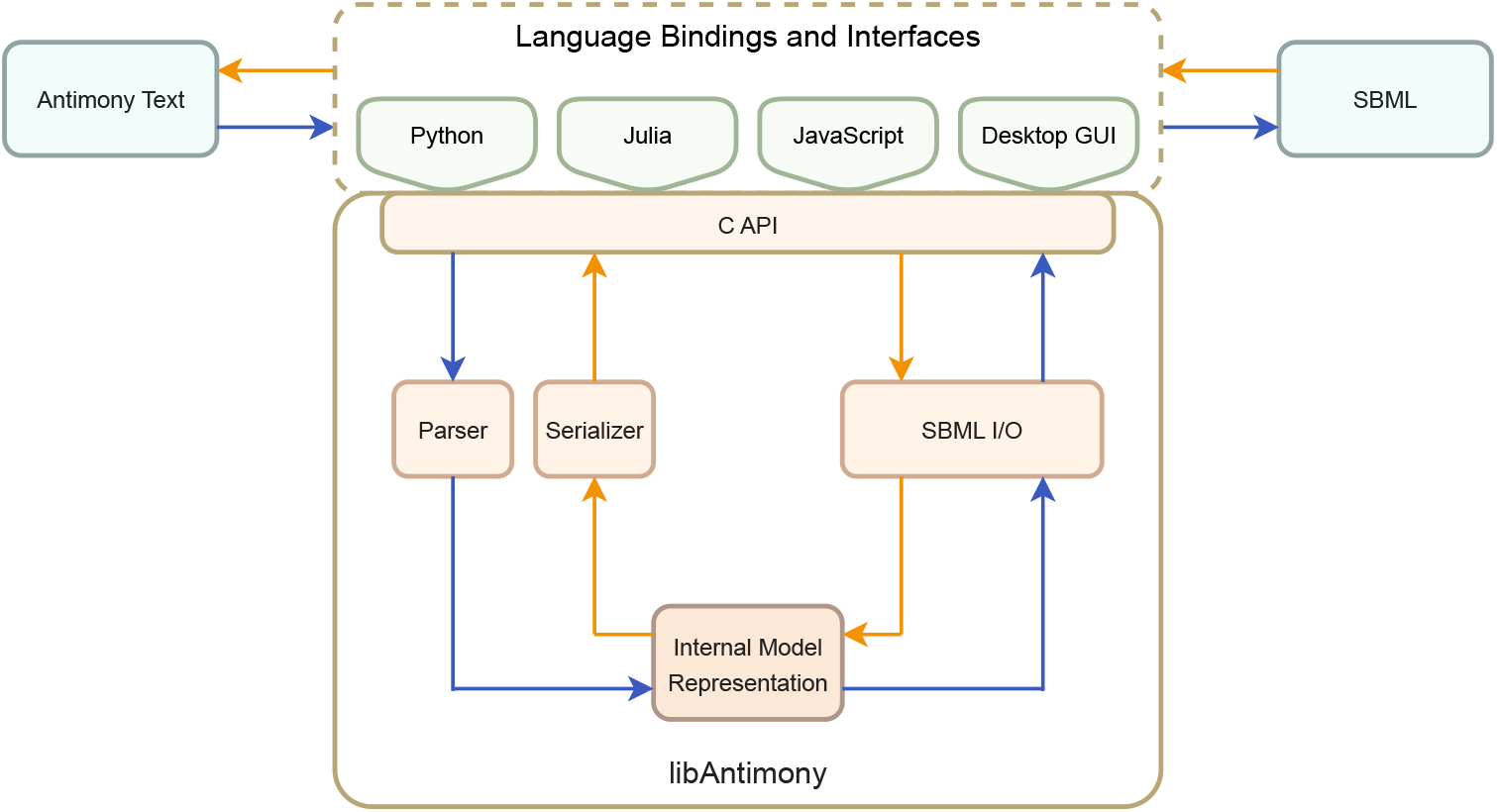
Antimony architecture and model translation pipelines. Blue arrows show Antimony-to-SBML pipeline: Antimony text enters through language bindings and the C API, is parsed by a Bison-generated parser into a hierarchical internal model representation, then mapped to an `SBMLDocument` via libSBML and written to an SBML (`.xml`) file. Orange arrows show the reverse SBML-to-Antimony pipeline: SBML files, provided through the language bindings and C API, are parsed by libSBML into an `SBMLDocument` object model, used to populate the same internal model representation, and serialized deterministically to Antimony syntax.

##### Internal Model Representation

At the core of Antimony’s architecture is a hierarchical internal model representation that bridges the Antimony syntax and the SBML format. This representation stores all model data translated from either format and ensures lossless conversion back to both.

##### Parser

The translation from the Antimony syntax to internal model representation is performed by a single-pass parser built with Bison [23], an established parser generator that generates parser logic from grammar specifications. Bison generates C++ code from a grammar file that is then incorporated into the libAntimony library to read the Antimony syntax line by line, conduct semantic validation, and construct the associated objects of each model element for the internal model representation.

##### Serializer

The reverse translation from the internal model representation to the Antimony syntax is performed by a single-pass serializer. It deterministically traverses the internal model representation, iterating over high-level objects and their constituent elements. Using this context and Antimony’s syntax rules, it reconstructs the corresponding Antimony syntax for each element.

##### SBML I/O

Antimony’s core statically links to libSBML [7], a mature and widely adopted library for constructing, querying, and serializing SBML documents, to handle translation between its internal model representation and the SBML format. Mirroring the parser/serializer pair used for the Antimony syntax, Antimony’s SBML I/O is organized as an importer and an exporter, implemented via libSBML. On import, Antimony loads an SBML model, traverses the resulting `SBMLDocument` object model, and populates its own internal model representation. On export, it constructs an `SBMLDocument` object, walks the internal model representation, and maps its elements to the corresponding components in the SBML structure. Additionally, to streamline the use of the SBML Layout and Render packages, Antimony’s core statically links to libSBMLNetwork [24], a lightweight library built on top of these packages that both surfaces their primitives and provides high-level utilities for model visualization.

##### C API

Antimony exposes its functionality through a deliberately minimal C API, centered on simple load and write operations for model import and export. The API has remained unchanged since the original release to ensure backward compatibility with existing workflows. This stable API enables most tasks to be performed with only a handful of functions, which makes its integration straightforward while still supporting the full range of Antimony’s functionality. It also serves as the foundation for language bindings and higher-level interfaces that interact with the Antimony core engine. A C API helps provide a cross-language interface.

##### Antimony-to-SBML Translation Pipeline

As shown by the blue arrows in Figure 1, this pipeline begins when Antimony text is passed through the external interfaces and routed via the C API into the core engine. Within the core, the process proceeds in three stages. First, the Antimony parser loads and processes the Antimony text, constructs the internal model representation, applies default values where needed, and reports any syntax or semantic errors. Second, the internal model representation is mapped to an `SBMLDocument` using libSBML. Third, the `SBMLDocument` is serialized to an SBML (`.xml`) file or string via libSBML writing functionality. The completed SBML output is then returned through the C API and the interfaces.

##### SBML-to-Antimony Translation Pipeline

The reverse path, shown by the orange arrows in Figure 1, begins when an SBML (`.xml`) file is provided through the external interfaces and passed to the core engine via the C API. libSBML parses the XML and constructs an `SBMLDocument` object model, which Antimony then traverses to populate its internal model representation. Once the representation is populated, the serializer walks through it and reconstructs the corresponding Antimony syntax as plain text. The generated Antimony text is then returned to the caller through the C API and the interfaces.

#### Dependencies

Antimony 3 is implemented in C++ and designed with minimal external dependencies to ensure easy compilation and deployment across platforms, while still leveraging established libraries for key functionality. These dependencies are listed below.

- **Bison**. Antimony uses Bison [23], a well-established GNU parser generator that turns grammar specifications into parsers, to define its parser logic and generate the corresponding C++ source file. The generated file is included in the source repository, so Bison is not required to build or install Antimony unless the grammar is modified and the parser needs to be regenerated.
- **libSBML**. Antimony statically links to libSBML [7], a widely adopted library for constructing, querying, validating, and serializing SBML models, and uses it to perform all SBML import/export operations. Static linking enables Antimony to be distributed as a self-contained library without requiring a separate libSBML installation.
- **libSBMLNetwork**. To support the SBML Layout and Render packages, Antimony statically links to libSBMLNetwork [24], a lightweight library built atop those packages that surfaces Layout and Render primitives and provides high-level utilities for model visualization. Similarly, because it is linked statically, no separate libSBMLNetwork installation is required at runtime.

#### Language Bindings and Interfaces

The modular architecture of Antimony 3, illustrated in Figure 1, enables its distribution across a broad range of environments through multiple interface modalities. At its foundation lies a C++ core library that exposes its functionality via a stable C API. This API serves as the basis for all language-specific bindings and front-end components. All other elements, including language bindings, graphical interfaces, and browser-based tools, are optional layers built atop this core and extend Antimony’s functionality into scripting environments, desktop applications, and web platforms. Supported distribution targets in Antimony 3 are described below.

- **C/C++ library**. The core distribution of Antimony 3 is a self-contained library that provides static and shared builds with C and C++ interfaces, enabling direct integration into native applications. The C interface also allows languages with C interoperability to access Antimony’s functionality via standard linking or foreign function interfaces.
- **Python bindings**. While earlier versions of Antimony relied on SWIG-generated wrappers [25], Antimony 3 adopts a lightweight `ctypes` [26] layer that is automatically generated from the C header. This approach eliminates the need for recompilation across Python versions. The Python bindings are distributed independently as pip-installable packages and are also bundled with Tellurium [17], a Python-based modeling environment for systems and synthetic biology, where Antimony serves as the primary language for defining and editing models within notebooks and scripts.
- **Julia bindings**. Similar to the Python `ctypes` layer, the Julia integration uses the language’s built-in `ccall` mechanism to invoke Antimony’s C API directly in Julia. This interface is released as a Julia package and provides support for seamlessly exposing Antimony’s functionality within the Julia environment.
- **Desktop GUI application**. For users who prefer a visual editing environment, Antimony provides a cross-platform desktop application (QtAntimony) built with the Qt framework. The interface features tabbed panes for simultaneously viewing and editing models in both Antimony syntax and SBML format. Real-time bidirectional translation, powered by libAntimony, enables users to switch seamlessly between textual and XML views while efficiently tracking modifications made to their models.
- **JavaScript/Web integration**. The Antimony core can be compiled to WebAssembly [27] using Emscripten [28], resulting in a lightweight JavaScript binding (`libantimony.js`) that runs entirely in the browser. This client-side configuration eliminates the need for server-side infrastructure and enables web applications to access Antimony’s functionality through direct JavaScript calls to its C API. A notable example of its adoption is MakeSBML [18], a single-page web application hosted statically on GitHub Pages and enables users to load, edit, and convert models between Antimony and SBML formats directly in the browser.

## Results

### Component and Model Annotations

Annotations are used to describe a model with references to common ontologies. We will distinguish annotations into two groups. One group is used to unambiguously describe the various biophysical components that make up a model. The second group is used to describe the overall aspects of a model. Both annotations make it possible to carry out more precise search, comparison, and reuse of models.

Component annotations are used to connect user-specific naming conventions for species, reactions, and parameters to standardized identifiers in well-characterized public databases such as ChEBI [29] and KEGG [30], or to chemical descriptors like SMILES [31].

Over the past decade, annotations have become increasingly central to modeling practices, driven both by the encouragement from BioModels curators [32, 33] to include rich semantic links and by the release of densely annotated models from the BiGG repository [34]. As researchers worked with these resources, they began to recognize their benefits and incorporate similar annotations into their own work. Despite this uptake, earlier versions of Antimony [11, 35] lacked native support for SBML annotations, which caused annotated SBML models translated to Antimony and back to lose those annotations.

To resolve this issue and address user requests, we now include annotation support in Antimony, allowing annotations to be added or edited directly in the Antimony model and preserved losslessly while round-tripping through the Antimony–SBML pipeline. In addition, to enhance the annotation capabilities of SBML models, we proposed and implemented an update to libSBML that extends the existing HTML support in human-readable notes to also include Markdown. This update enables Antimony annotations to include the HTML notes as Markdown, and ensures that formatting is retained when converting between Antimony and SBML.

Antimony 3 provides a compact, declarative syntax to attach annotations to any model element with an id, including the model itself. The core annotation form is `[identifier] [qualifier] “[URI]”`, where the `identifier` is the id of the model element being annotated, the `qualifier` is a controlled keyword, and the `“URI”` points to an external resource, often provided through `Identifiers.org` [36]. Human-readable annotations in Markdown can also be attached to elements using the `notes` quantifier.

Listing 1 illustrates an Antimony excerpt from `BIOMD0000000004` [37] that uses all three types of annotation. At the element level, compartments, species, and reactions are linked to standard ontologies and databases (e.g., GO [38], InterPro [39], UniProt [40]). At the model level, annotations capture identity, provenance, cross-references, and contributor information and helps document details about the authors and provenance of the model and provide links to related repositories, archived releases, and associated publications [41]. The model notes block provides human-readable context and additional information about the model in Markdown format.

### Flux Balance Constraints

Constraint-based modeling is an approach based solely on the stoichiometric structure of a model and takes no account of the kinetic and regulatory aspects of a metabolic network. Historically the technique have been used primarily to model genome-scale models, and often employs linear programming coupled to an objective function as well as additional constraints and bounds [13] to estimate steady state fluxes.

SBML Level 2 [42] had no official way to encode constraint-based models, so software tools adopted pragmatic approaches to work around this limitation. For example, the COBRA Toolbox [43] used SBML’s `Species` and `Reaction` classes to represent the reaction network, while FBA-specific components were handled through ad hoc conventions, most notably by relying on shared naming conventions for local parameter `id`’s across tools. As methods and applications of genome-scale constraint-based models expanded, the need for a standard to define, exchange, and annotate these models became essential. In response, the Flux Balance Constraints (FBC) package [13] was introduced to extend SBML Level 3 Core with the elements required to encode such models.

Antimony 3 introduces a human-readable syntax for defining constraint-based models. Objectives are specified using the keywords `maximize` or `minimize`, followed by a linear combination of reaction `id`s (e.g., `maximize R1 + R2`). Additionally, flux bounds are defined directly on reaction `id`s using inequalities to set limits or equalities to fix a flux (e.g., `0 <= R1 <= 1000`).

Listing 2 illustrates how a constraint-based model is declared in Antimony, using an excerpt from BIOMD0000001046 [44], which defines a model of the mycolic-acid pathway. The excerpt highlights the three core ingredients of a constraint-based model in Antimony: (i) a stoichiometric reaction network, here represented by a set of export (exchange) reactions for multiple mycolate classes that are combined into a biomass pseudo-reaction using their stoichiometric weights; (ii) an optimization objective, declared to maximize the flux of this biomass reaction; and (iii) flux bounds, specified through upper and lower inequalities applied to reaction identifiers to constrain allowable reaction fluxes.

### Distributions

Biological models often rely on parameters and initial conditions that are not fixed values but are instead uncertain or inherently stochastic across different contexts. For example, when modeling heterogeneous cell populations, metabolite levels may vary from cell to cell, so simulating a batch requires sampling from a distribution rather than assigning a single fixed value. Similarly, in pharmacological applications such as pharmacokinetic/pharmacodynamic (PK/PD) models, variability in patient physiology is best represented by running repeated simulations that draw parameter values from distributions. In addition, some biological processes are inherently stochastic. For instance, in gene regulatory network models, a random trigger can be used to represent transcriptional bursts or stochastic regulatory events.

**Listing 1.**
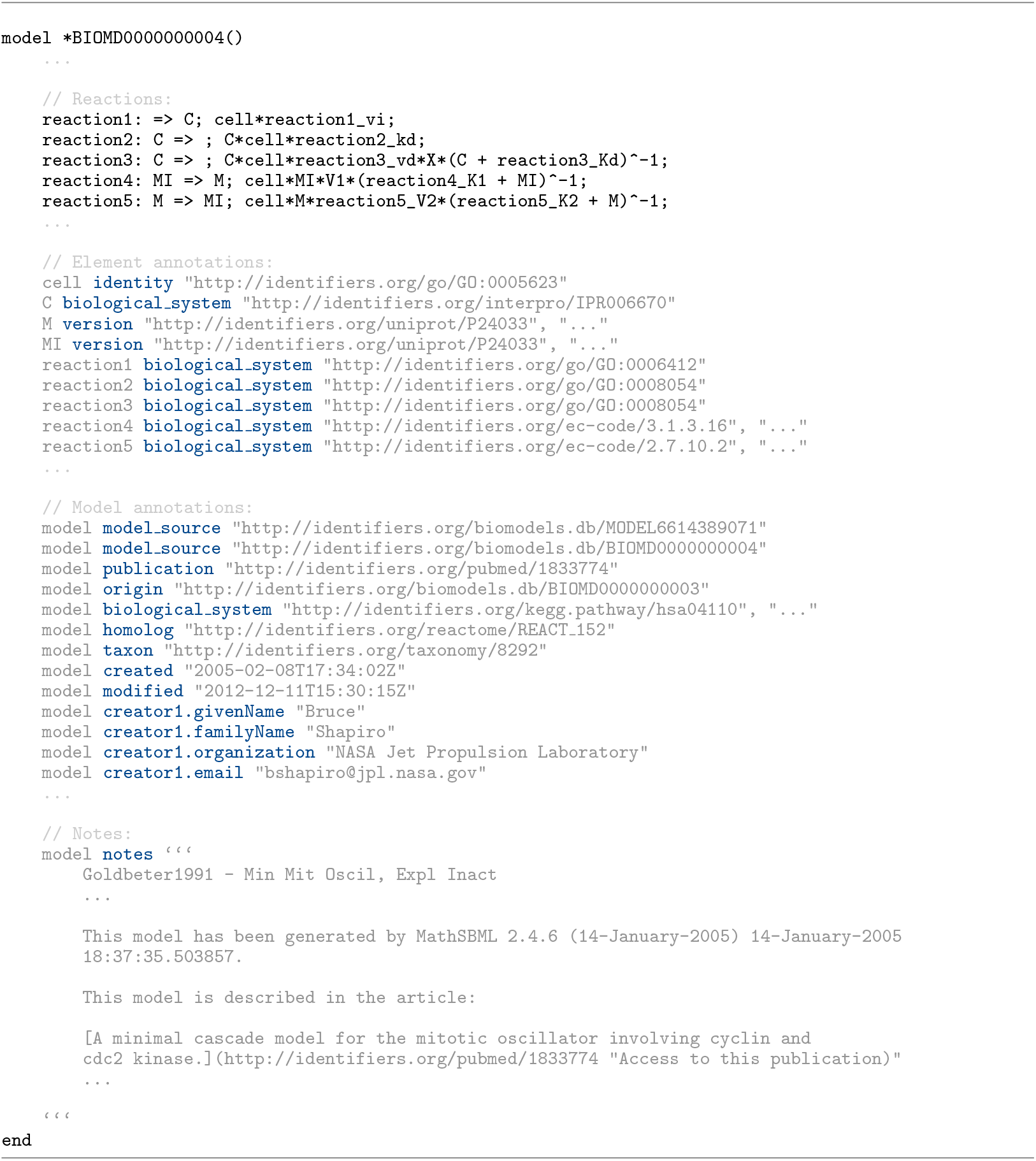
Antimony excerpt from BIOMD0000000004 [37] illustrating all three types of annotation: element-level links to standard ontologies and databases, model-level metadata and provenance, and a Markdown notes block providing additional inofromation about the model

**Listing 2.**
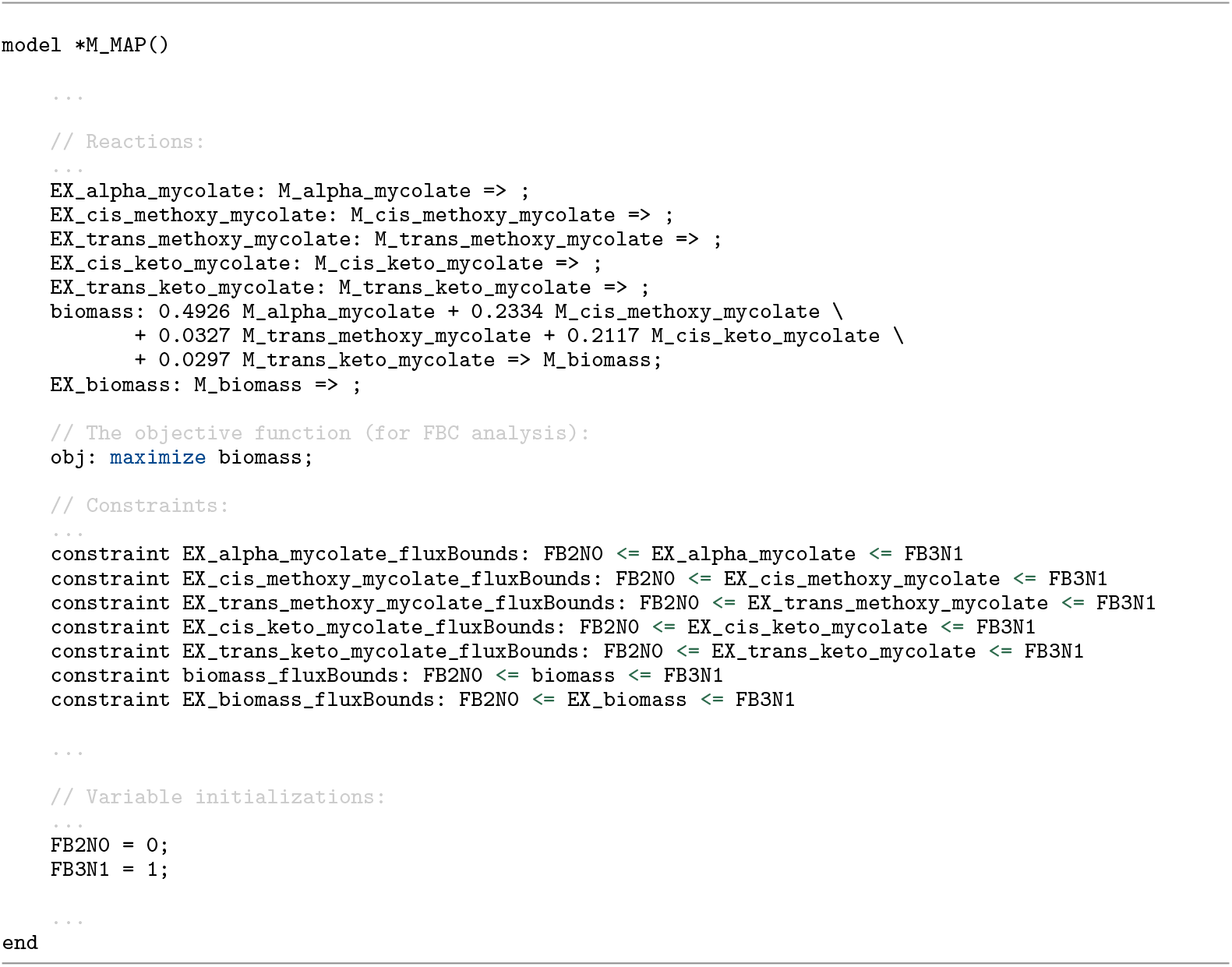
Antimony excerpt from BIOMD0000001046 [44] illustrating the declaration of a constraint-based model using Flux Balance Constraints (FBC). The model includes export (exchange) reactions aggregated into a biomass pseudo-reaction, an optimization objective that maximizes biomass flux, and reaction flux bounds which are specified as upper and lower inequalities.

SBML Level 3 Core had no explicit way to encode uncertain or random values within a model. In practice, the common approach among modelers was to generate batches of simulations with sampled values in external scripts or simulator settings, leaving SBML files with only nominal values and no explicit record of uncertainty.

The Distributions package [14] was introduced to provide a standard way of encoding uncertainty and stochasticity directly within SBML models. It adds two complementary capabilities to the standard. First, it extends SBML Math with functions to draw samples from probability distributions such as `normal`, `uniform`, or `lognormal`. These functions can be used within eligible mathematical expressions in the model, and, when invoked, generate values from the specified distribution. Second, it provides a structured approach for documenting uncertainty in element values through statistics such as mean, standard deviation, and variance, which can be attached directly to SBML elements and preserve variability information alongside their nominal values.

Antimony 3 integrates the capabilities of the SBML Distributions package into its human-readable syntax, which allows modelers to introduce uncertainty and stochastic behavior straightforwardly into their models. Parameters and initial values can be specified in Antimony 3 by drawing samples from probability distributions, using the same set of functions defined in the Distributions extension of SBML. Listing 3 shows a PK/PD model [14] in Antimony format where the `central` compartment volume and kinetic parameters, `ka` and `ke`, are initialized by draws from a `lognormal` distribution. Once the initializers are evaluated, these values are sampled from the distribution, so each simulation run may begin with different parameter and compartment values while the overall model structure remains unchanged.

**Listing 3.**
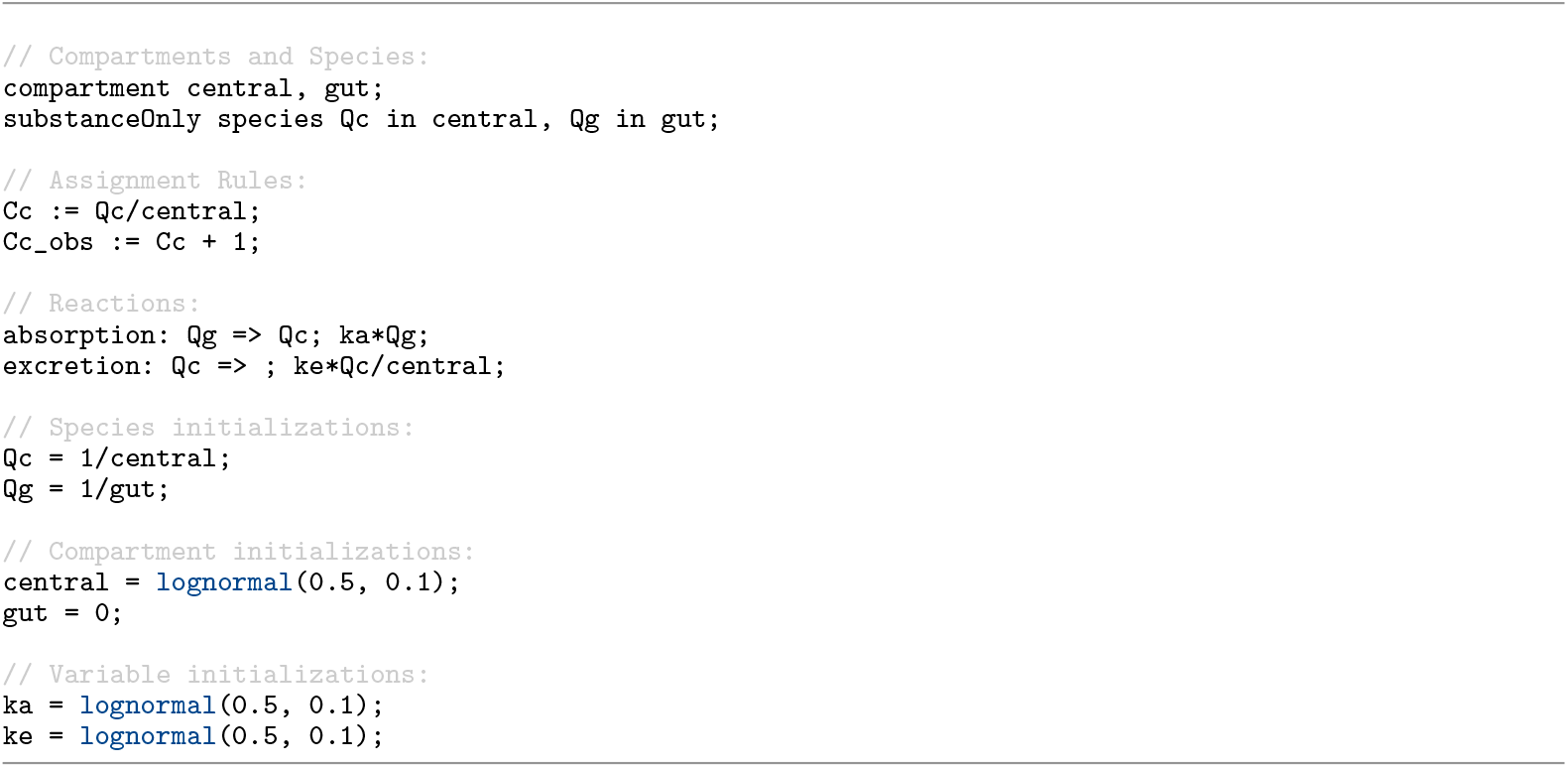
A PK/PD model [14] in Antimony format demonstrating initialization from probability distributions. The `central` compartment volume and kinetic parameters (`ka`, `ke`) are initialized by draws from a `lognormal` distribution.

Probability distributions can also be used in discrete events, either in the trigger to model random event times or in the assignment to reset a parameter or component value to a newly sampled value when the condition is met. Listing 4 shows a model [14] in Antimony format where two simultaneous events, `E0` and `E1`, are triggered when the condition `time > 2` and `x < 1` is met, and are ordered based on their `priority` values sampled from `uniform(0,1)` and `uniform(0,2)`. In this example, because the priority of `E1` is sampled from a wider distribution, it has the higher priority in about `75%` of runs and fires first, thus setting `x = 5` and canceling `E0`. In the remaining `25%`, `E0` executes first and sets `x = 3`.

**Listing 4.**
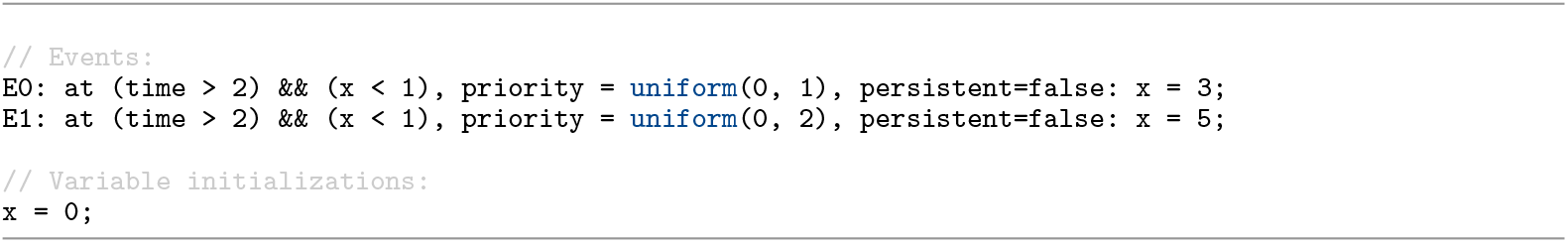
Antimony events using probability distributions in their priorities [14]. Two simultaneous events, `E0` and `E1`, are triggered when `time > 2` and `x < 1`. Their priorities are sampled from `uniform(0,1)` and `uniform(0,2)`; in about `75%` of runs, `E1` has the higher priority, fires first, and sets `x = 5`, which cancels `E0`. In the remaining `25%`, `E0` fires first and sets `x = 3`.

In addition, Antimony 3 supports the attachment of uncertainty metadata to model elements. The same statistics defined in the Distributions extension of SBML can be associated with any symbol (e.g., parameters, species, compartments, or reactions) in the Antimony format. As an example, Listing 5 shows a model [14] in Antimony format where dot notation is used to attach uncertainty statistics to model elements (standard deviations for species and confidence intervals for parameters). It should be noted that, consistent with the Distributions specification, these annotations serve only to document the statistical context of values and do not affect simulation dynamics.

**Listing 5.**
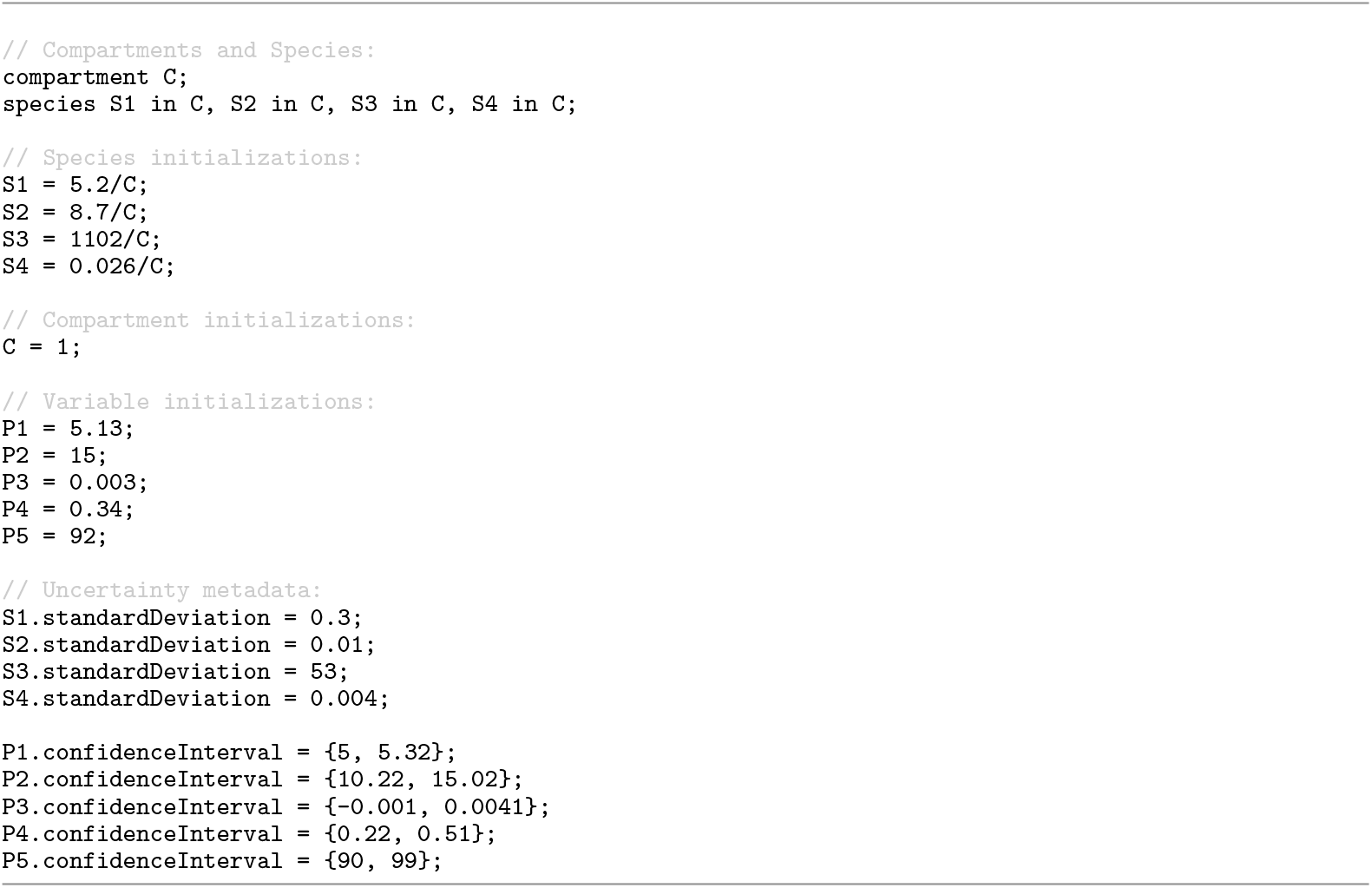
Uncertainty metadata attached to model elements via Antimony’s dot notation [14]. Species (`S1-S4`) are annotated with `standardDeviation` values and parameters (`P1-P5`) with `confidenceInterval` ranges.

### Layout and Render

Visualization of biological models, by graphically depicting network topology and overlaying simulation or experimental results, provides a powerful means of enhancing the understanding of complex models and their dynamic behavior. This approach makes intricate relationships within a model more comprehensible and helps researchers effectively communicate their findings and insights to a broader scientific community [24]. However, the absence of a widely accepted visualization standard for biological models can lead to a variety of challenges, including limited interoperability, cumbersome data exchange, and poor reproducibility of visualization data across different tools and platforms.

To address these challenges, the SBML Level 3 Layout [15] and Render [16] packages were introduced as community standards for encoding model visualization alongside the SBML model. The Layout package captures the structural arrangement of model elements, while the Render package defines their graphical style. By encoding visualization data directly within SBML, these packages ensure that model diagrams remain reproducible and portable across tools, amd their visual glyphs faithfully linked to the underlying model elements. No additional effort is required to maintain separate files for the model and its visualization.

**Listing 6.**
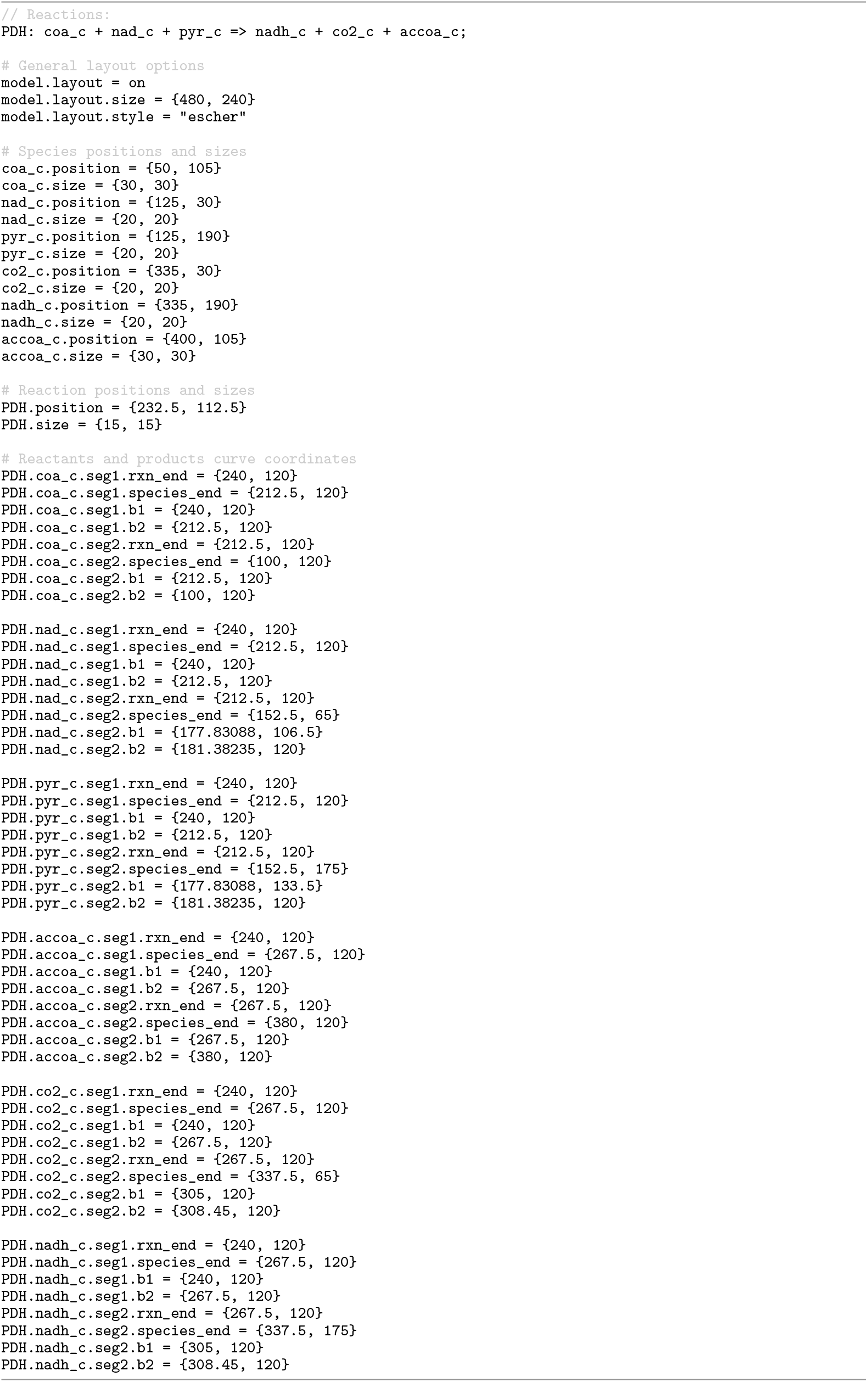
Antimony excerpt encoding visualization information for a minimal pyruvate dehydrogenase reaction model using the SBML Layout and Render packages. The listing specifies species and reaction centroid coordinates, as well as reaction curve segment coordinates, and applies an Escher-style visual template [45]. The rendered output generated from these specifications is shown in Fig 2.

Antimony 3 introduces native support for the SBML Layout and Render packages, extending its human-readable syntax to include visualization details alongside model definition. In practice, most SBML models do not include visualization details encoded in the Layout and Render specifications. To address this gap, Antimony 3 leverages libSBMLNetwork’s auto-layout feature [24] to generate an initial diagram that provides a practical starting point for visualization. This means that the only necessary Antimony directive for creating and saving a visual representation of a model is the line `model.layout = on`. Details such as the position or color of particular elements may be added as needed. Whether the diagram is auto-generated or already present, its visualization data is imported and represented in Antimony syntax for inspection and modification. Using its concise dot-notation, Antimony 3 then enables modelers to customize visualizations, from applying predefined styles for rapid theming to making fine-grained adjustments to specific elements, all within the Antimony format.

To demonstrate Antimony support for model visualization through the SBML Layout and Render packages, Listing 6 presents a minimal model of the pyruvate dehydrogenase reaction configured with the Escher-style template [45]. The Antimony model in this listing integrates the definition of the model with visualization details through specifying the positions and sizes of the species and the reaction centroid, setting the reaction curve coordinates, and applying a visual template in Escher style. This Antimony model can be translated into SBML, where its visualization information is encoded in the Layout and Render packages format. The resulting SBML model can then be visualized using tools that support these packages, such as SBMLNetwork [24] (which we used to generate the diagram shown in Figure 2).

**Fig 2.**
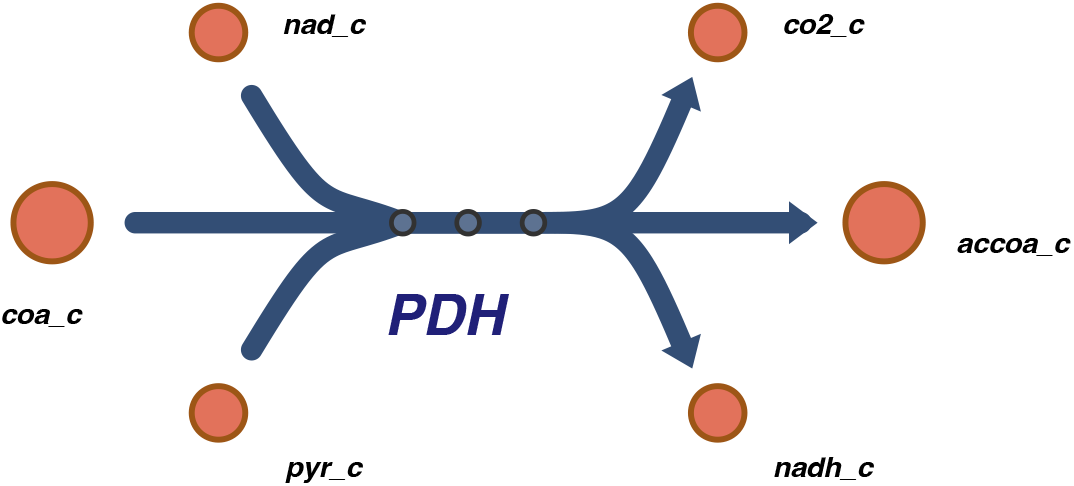
Rendered visualization of the pyruvate dehydrogenase reaction defined in Listing 6. Drawn by SBMLNetwork [24] using visualization data specified in the Antimony model. As illustrated in Listing 6, Species positions and Reaction centroid coordinates are defined explicitly, reaction curves are specified via segment coordinates, and the Escher style template [45] determines the overall appearance.

### rateOf

In a dynamical analysis of a model, it is often necessary to retrieve or reference the rate of change, with respect to time, of an entity. For example, in regulatory network models, a common strategy is to monitor the change of a key readout to quantify the strength of a transient pulse and to trigger interventions only during periods when that readout is rising or falling sharply [46]. In multicellular systems models, the rate of change of a supporting cell population can be referenced to determine when dependent populations should be penalized or protected [47]. In compartmental epidemic models, the rate of change of the infected population can be used to automatically identify the peak of an outbreak, which provides a basis for structured evaluation of alternative scenarios [48].

SBML originally provided no standard way to retrieve or reference a rate of change inside a model equation. Modelers instead relied on manual workarounds to approximate derivative values, or turned to tool-specific constructs such as COPASI’s custom annotations method for representing rates of change in SBML [6]. SBML Level 3 Version 2 resolves this limitation by introducing the `rateOf`symbol, a built-in construct that allows equations to reference the time derivative of eligible symbols directly within SBML Math [12]. This provides a standard way to access rates of change in equations and eliminates the need for ad hoc conventions or tool-specific extensions.

Antimony 3 exposes this capability through the function call `rateOf(<id>)`, where the argument must be the identifier of an eligible model symbol. This allows the rate of change of that symbol to be inserted directly into equations such as reaction rate laws, assignment rules, as well as event triggers and assignments. To demonstrate this capability, Listing 7 shows an excerpt from an Antimony model of brain tumour, `BIOMD0000000775` [47], where the rate of change of the glial cell concentration (`G`) is captured using `rateOf`, passed through a Heaviside gate (`H`), and reused in the rate law of neural population reaction (`n increase`). As a result, the contribution of glial cell concentration to this reaction is effective only when it is decreasing, with a magnitude proportional to the rate of its decrease.

**Listing 7.**
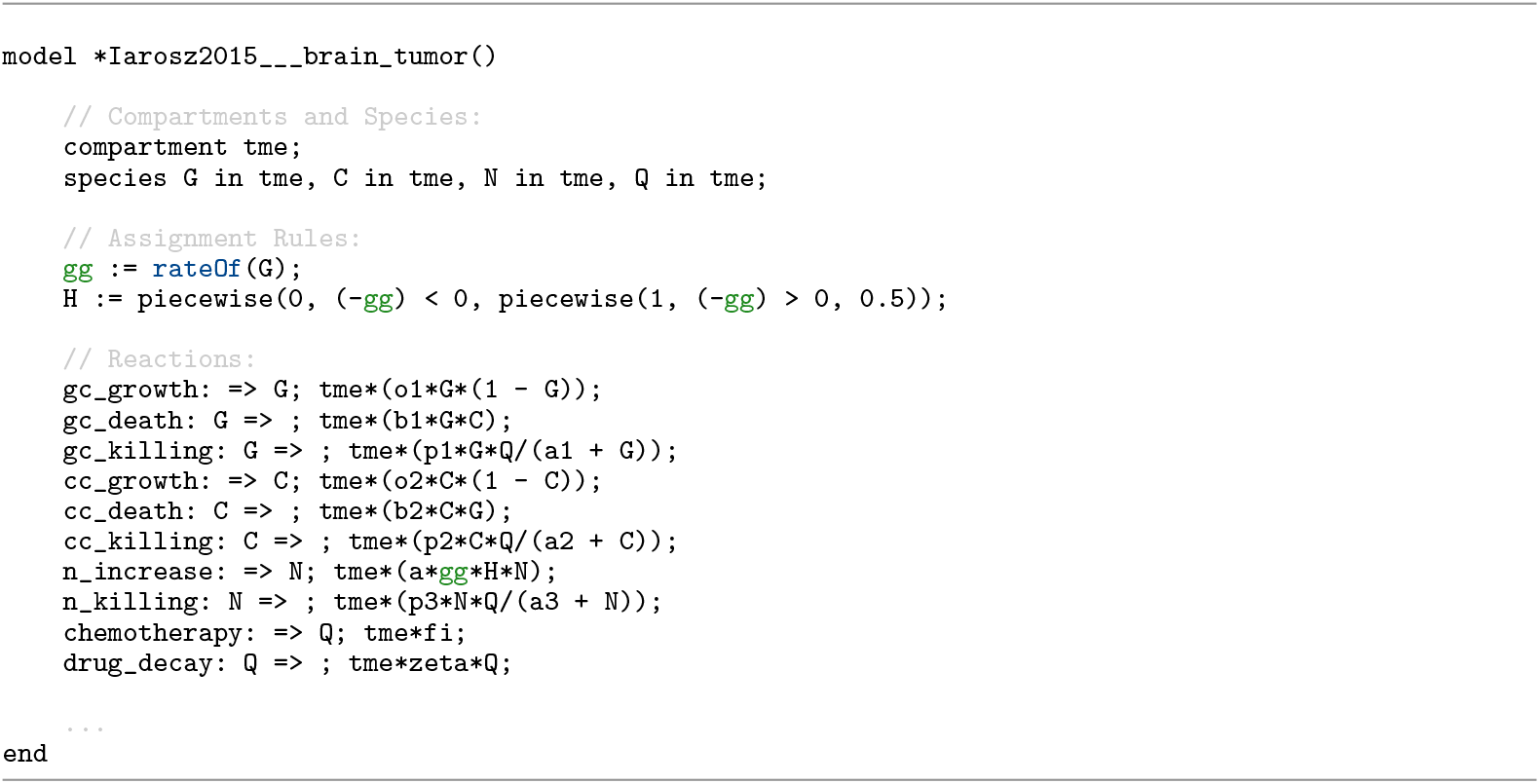
Antimony excerpt from BIOMD0000000775 [47] showing the use of the `rateOf` construct. The rate of change of glial cell concentration (`G`) is retrieved, mapped through a Heaviside gate (`H`), and reused in the neural population reaction (`n increase`) rate law, contributing only while `G` is declining and with magnitude proportional to the rate of that decline.

As another example, the use of assignment rules with the `rateOf` provides a way to model a classical proportional-integral-differential (PID) control for a biological system. Listing 8 displays such an implementation where `rateOf` is used to implement the differential control. Fig. 3 shows that in a simulation with this differential control (the last frame in the figure), the system is robust to a sinusoidal disturbance.

**Listing 8.**
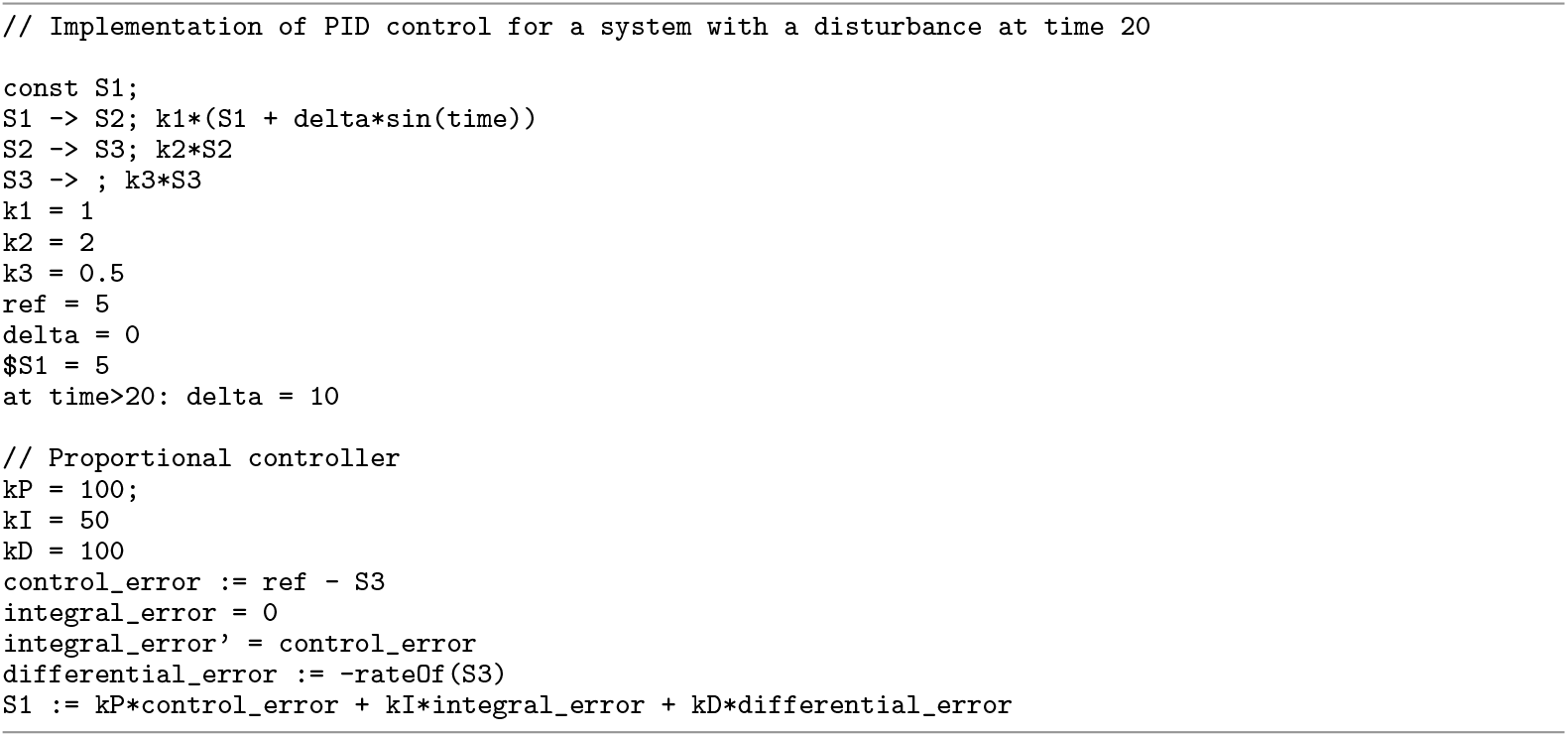
Implementation of Proportional-Integral-Differential (PID) control for a system with a disruption at time 20. The `rateOf` construct is used to define `differential error`.

**Fig 3.**
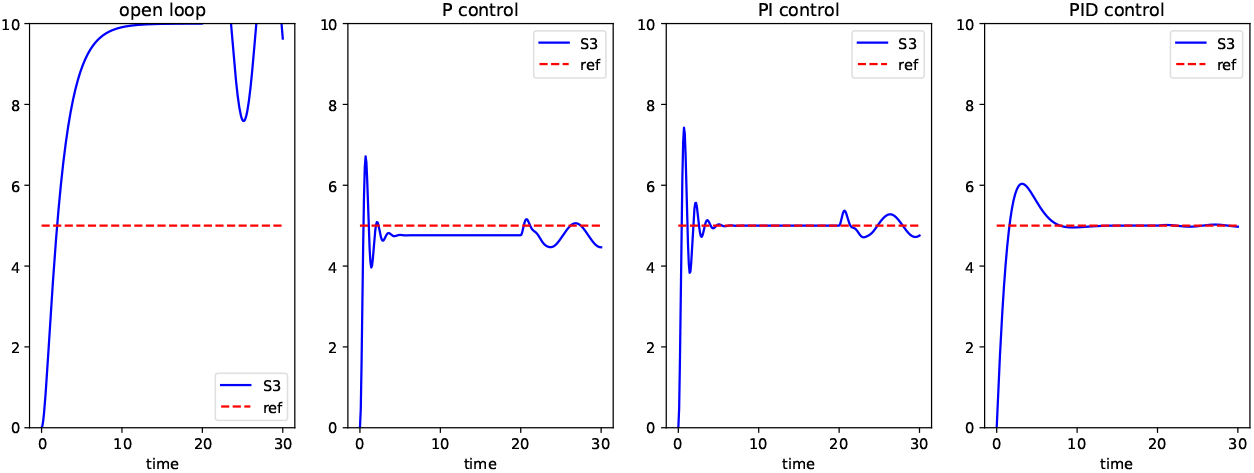
Timecourse produced by the PID controller. The figure displays the time courses for different controller designs of the model in Listing 8 by the variables kP (proportional control), kI (integral control), and kD (differential control). The objective is to maintain the output (S3) at the reference of 5 in the presence of a sinusoidal disturbance at time 20. In “open loop”, kP = kI = kD = 0, and the objective is not achieved. In proportional control, kI = kD = 0, and we are close to the reference but the disturbance is still visible. In proportional-integral control, kD = 0 and we maintain the reference, but not during the disturbance. In proportional-integral-differential control (which uses the `rateOf` construct in the model), the reference is maintained in the presence of the disturbance.

### Algebraic Rules

SBML *Rules* are used to define the values of variables in a model, their relationships, and their behavior over time. SBML distinguishes three types of rules: *Assignment* rules, which set a variable’s value as a function of others, *x* = *f*(*V*); *Algebraic* rules, which impose constraints of the form 0 = *f*(*W*) that must hold throughout a simulation; and *Rate* rules, which specify how a variable changes over time, 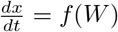 (here, *V* denotes symbols that exclude *x*, while *W* denotes symbols that may include *x*) [12].

Although assignment and algebraic rules share a similar mathematical structure, SBML distinguishes them for practical reasons. Assignment rules serve as an explicit subclass of algebraic constraints, making them well suited for defining intermediate quantities in numerical methods. In addition, certain numerical analyses can only be performed on models without algebraic rules, which reinforces the need to keep the categories distinct. At the same time, many simulators do not support solving unconstrained algebraic equations. As a result, expressing these relationships as assignment rules broadens the range of tools that can execute the model [12].

Antimony 3 introduces native support for SBML algebraic rules, where the syntax of an algebraic rule is defined as `[ruleId:] 0 = <algebraic-expression>`. To demonstrate how algebraic rules are declared, Listing 9 provides an Antimony excerpt from a model in the SBML Level 3 Version 2 specification [12], which contains two algebraic rules that cannot be expressed as assignment rules. The model describes a quasi-steady-state approximation of the enzymatic reaction:

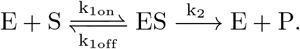

For this reaction, assuming that the intermediate complex is at steady state, 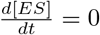, and that the total enzyme concentration is conserved, [*E*] + [*ES*] = [*E*_total_], leads to an algebraic rule of the form 0 = *f* ([*ES*]), in the model.

**Listing 9.**
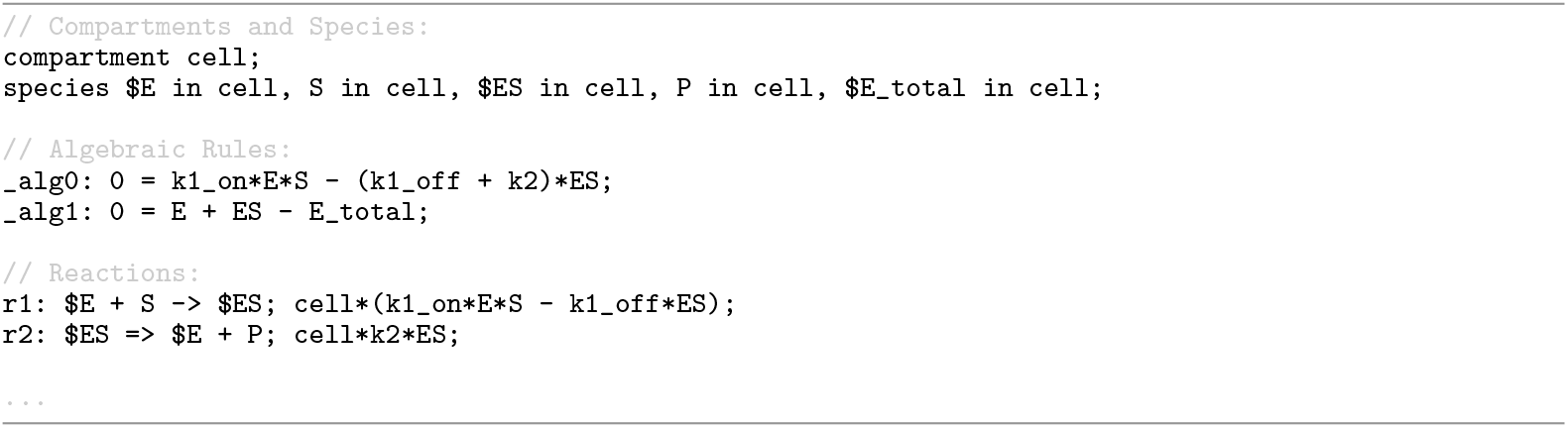
Antimony representation of a model from the SBML Level 3 Version 2 specification [12], illustrating native support for algebraic rules in a quasi-steady-state enzymatic reaction model.

### substanceOnly

In SBML, a species identifier can represent either an amount (substance units) or a concentration (amount divided by compartment size) when it appears in mathematical expressions. While it is always possible to convert between the two by multiplying or dividing by the compartment size, making the intended interpretation explicit keeps mathematical expressions simpler and avoids ambiguity. However, this interpretation is not determined by how the species’ initial value is set. SBML allows initial values to be provided in different ways (e.g., `initialAmount` and `initialConcentration`), but they are all optional attributes. In addition, in many SBML models initial values are unknown, supplied from an external source, or determined by an initial assignment [49]. Therefore, SBML makes the interpretation of a species identifier explicit using the `hasOnlySubstanceUnits` attribute. If `hasOnlySubstanceUnits = true`, the species identifier in expressions represents an amount; if it is `false`, the identifier represents a concentration.

Earlier versions of Antimony [11, 35] treated all species identifiers as concentrations. While this worked well for many kinetic models, it made it difficult to represent models that are intentionally built in terms of species amounts. This was especially the case when modelers wanted to always think in terms of amounts, particularity because reactions’ kinetic laws in SBML always change the amounts of species, not their concentrations.

To preserve SBML semantics and support these types of models, Antimony 3 introduces the `substanceOnly` qualifier for species declarations as a human-readable representation of SBML’s `hasOnlySubstanceUnits = true`. Species identifiers can now be made to represent amounts in mathematical expressions using the declaration `substanceOnly <id1>, <id2>,` …. In addition, Antimony’s initialization grammar is updated accordingly to reflect this interpretation. A statement such as `S1 = 10` previously always meant “initial concentration.” Now, if `S1` is declared `substanceOnly`, the same statement means “initial amount.”

### Named stoichiometries

In an SBML Level 3 reaction, the stoichiometry of an individual reactant or product can be given an id, and that id can be used in mathematical equations (where it represents the value of that stoichiometry), and can be the target of initial assignments or rules. For some models, the precise mechanism being modeled is not known, so different stoichiometry values need to be set to compare with experimental data. An example of its usage is in a model for the Field-Noyes Model of a Belousov reaction [50], where the ‘stoichiometric factor’ might be 1, 1/4, or smaller, depending on the behavior of Ce(rV), bromomalonic acid, and malonic acid. In Antimony 3, it is now possible to construct a reaction with an id instead of an explicit stoichiometry, and that id can then be used elsewhere in the model. Listing 10 illustrates this with a simple model where the stoichiometry of a ligand `L` is given the id `n`, which is then used in the reaction’s rate equation, and set to be equal to 3. This model is then used (Fig 4) to illustrate the effects of cooperativity by analyzing steady state values at different values of `n`.

**Listing 10.**
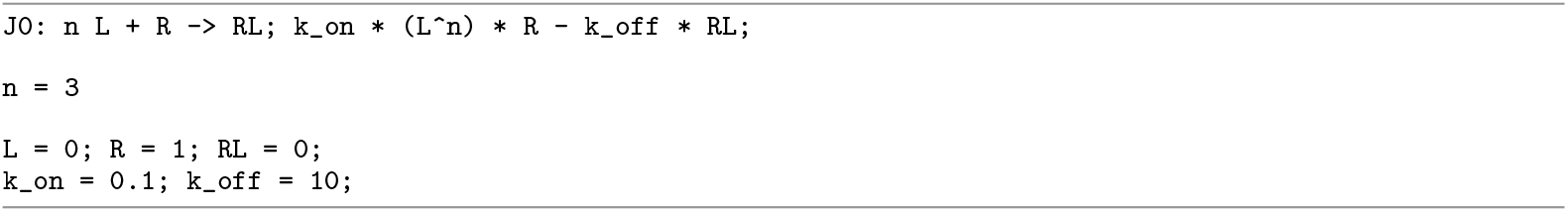
Antimony representation of a model with a named stoichiometry (`n`).

**Fig 4.**
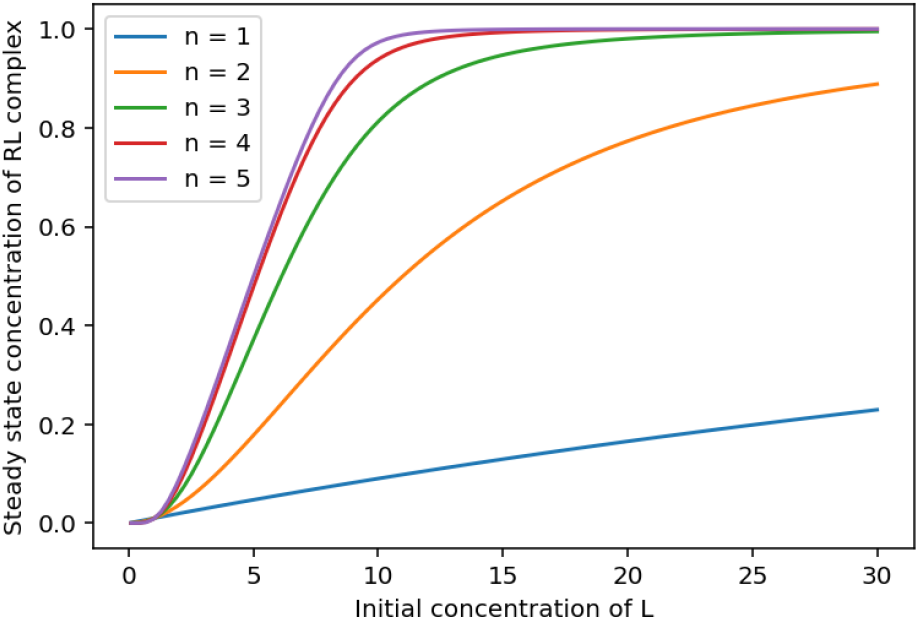
Steady state analysis of differential stoichiometric analysis. An analysis of the model in Listing 10 plotting the effects of different initial values of the ligand `L` on the final concentration of the complex `RL` at steady state, for different values of `n` (the stoichiometry of `L`).

## Discussion

Antimony was originally introduced to address a fundamental challenge in systems biology modeling: how to make the increasingly sophisticated capabilities of modern modeling standards accessible to domain scientists without compromising standards compliance, interoperability, and reproducibility. Traditional solutions to this problem have tended to favor one side of the balance. Graphical tools prioritize intuitive model construction and accessibility but often struggle to scale to complex models or to maintain strict standards compliance. At the other end, programmatic libraries expose the full breadth of standards but demand substantial programming expertise and time investment from users. Domain-specific languages emerged as a middle ground by offering more concise, human-friendly abstractions, yet most have remained tied to particular host languages or have provided only partial coverage of evolving standards. Antimony stands out in this space by offering a language-agnostic, plain-text syntax that round-trips to SBML without loss, which offers both enhanced human readability and full SBML compliance.

Since the first Antimony release, SBML has matured to Level 3 Core and has been extended with a rich ecosystem of packages. This has introduced constructs that the original Antimony syntax could neither express nor preserve. In Antimony 3, we consolidate successive upgrades, implemented to keep pace with the evolution of SBML, into a single, comprehensive release: the language now covers SBML Level 3 Core, with broader support for advanced modeling features, together with the Flux Balance Constraints, Distributions, Layout, and Render packages, all while maintaining Antimony’s established syntax and ensuring backward compatibility with existing models.

The support for SBML Level 3 Core and its packages in Antimony 3 enables unified modeling workflows that were previously fragmented across multiple tools and formats. Native annotation capabilities address a long-standing limitation by allowing semantic metadata to be defined directly within model description, which prevents the loss of provenance and biological context during round-trip translations. Support for the Flux Balance Constraints (FBC) package enables constraint-based analysis to be expressed directly in Antimony 3, where objectives and flux bounds can be specified natively rather than through ad hoc scripts or tool-specific conventions. The incorporation of Distributions package in Antimony 3 makes uncertainty and stochastic behavior part of its syntax: parameters, initial conditions, and event constructs can be sampled from probability distributions, and uncertainty statistics such as means, standard deviations, and variance can be attached directly to symbols as metadata. Native support for the Layout and Render packages enables Antimony 3 to embed visualization details directly within model definitions. Diagrams import losslessly or auto-generate when absent and their details can be inspected or customized entirely within the Antimony format. Finally, support for particular SBML Level 3 Core elements enables modelers to seamlessly incorporate richer mathematical formulations into models. The `rateOf` construct provides standardized access to time derivatives in equations, the algebraic rules allow conservation assumptions to be encoded explicitly, and `substanceOnly` explicitly specifies whether species identifiers are treated as amounts or concentrations in the model’s mathematical expressions.

The collective impact of these capabilities becomes increasingly significant when considering complex modeling scenarios that require multiple advanced features simultaneously. Consider a genome-scale metabolic model of a cell population under varying environmental conditions: such a model might incorporate flux balance constraints to optimize system-level objectives, probability distributions to represent heterogeneity in metabolite concentrations, `rateOf` constructs to couple growth dynamics with flux variations, comprehensive annotations that contextualize reactions biologically by linking them to reference databases such as KEGG and BiGG, and visualization specifications for defining publication-ready pathway diagrams. Previously, constructing such a model would have required coordinating multiple tools, formats, and external scripts, with significant risk of losing crucial information during format conversions and making reproducibility difficult to achieve. With Antimony 3, this entire specification, ranging from mathematical formulation through semantic annotation to visual presentation, can be captured in a single, human-readable text file that remains accessible to domain scientists while maintaining full SBML compliance.

In addition to its advanced modeling capabilities, Antimony 3 benefits from a broad distribution strategy that supports its adoption across diverse user communities. Python users can install its dedicated package directly or access it through the established Tellurium environment. Julia users have access to a Julia package, while browser-based platforms can integrate its WebAssembly build for client-side execution without relying on server infrastructure [18]. For users who prefer graphical interfaces, the QtAntimony desktop application offers real-time, bidirectional translation between Antimony and SBML. By making Antimony 3 available within researchers’ preferred environments, the project lowers the barrier to adopting the SBML Level 3 features and promotes the broader use of these capabilities within the system biology community.

## Availability and Future Directions

Antimony 3 is open-source software distributed under the BSD 3-Clause License, with its source code available on GitHub at https://github.com/sys-bio/antimony. It supports multiple platforms and environments.

Native C/C++ distributions are provided as both static and shared libraries via the release page, and can be linked using a stable C API. For Python users, Antimony can be installed via PyPI using `pip install antimony`, or accessed as part of the Tellurium modeling platform (https://github.com/sys-bio/tellurium), which is installable with `pip install tellurium`. Julia users can use Antimony through the RoadRunner.jl package (https://github.com/sys-bio/RoadRunner.jl), installable via `add RoadRunner` in the Julia package manager.

JavaScript and WebAssembly bindings are available through the libantimonyjs project (https://github.com/sys-bio/libantimonyjs) release page, supporting browser-based applications. An online demonstration is hosted at MakeSBML (https://sys-bio.github.io/makesbml), a web app that converts between Antimony and SBML formats directly in the browser. Additionally, a cross-platform desktop application, QtAntimony, provides a graphical interface for converting between Antimony and SBML formats. Its precompiled binaries for Windows, macOS, and Linux are available on the GitHub releases page.

Antimony’s comprehensive documentation, including a detailed Antimony language reference, tutorials, and example models, is available at https://tellurium.readthedocs.io/en/latest/antimony.html.

As we continue to develop Antimony, our primary goals are to continue to cover new areas of modeling, particularly those covered by SBML Level 3 packages. Qualitative modeling (models where elements have explicit or Boolean levels instead of having continuous numerical values) has a robust community of modelers [51], and has generally settled on SBML with the ‘Qualitative Modeling’ package [52] as a standard format for exchanging models between tools. Rule-based modeling (where the interactions of types of reactions, species, and compartments are defined instead of explicitly modeling each individual one) similarly has many users, particularly in systems where the same sort of reactions occur in multiple compartments and/or across multiple variations. Model exchange is more rare, but when present, does use the ‘Multistate, Multicomponent and Multicompartment Species’ package [53]. A simulator-agnostic model creator like Antimony could be useful for encouraging model exchange in this domain. As other modeling domains adopt SBML as a model exchange format, Antimony development may pursue those domains as well.

Antimony has also proven to be an ideal target for creating biological models with the assistance of language models. The format is concise, formatting is light, and a single line of Antimony is sufficient to express a single concept in a model, making it an effective format for a variety of modeling approaches [54–57]. As language models and biological modeling approaches continue to evolve, we plan to further maintain and enhance Antimony’s suitability for language model–assisted model creation.

## Acknowledgments

HMS, JLH, LS, and AH acknowledge support from the Center for Reproducible Biomedical Modeling, which is funded by the National Institutes of Health (NIH) under grant number P41EB023912.

